# What causes different coronal curve patterns in idiopathic scoliosis?

**DOI:** 10.1101/2020.01.21.913707

**Authors:** Saba Pasha

## Abstract

**Background:** Adolescent idiopathic scoliosis (AIS) is a three-dimensional deformity of the spinal column in otherwise healthy adolescents. The underlying mechanisms associated with the spinal deformity development have been explored which delineated the role of the sagittal curvature of the spine. The patterns of the spinal deformity vary between the AIS patients as shown in several classification systems. It remains to further investigate how variations in sagittal profiles result in different coronal plane deformities in AIS and how these deformation patterns are intrinsically different.

**Methods:** A total of 71 Lenke 1 and 52 Lenke 5 AIS patients were included retrospectively. The 3D models of the spine were generated from biplanar radiographs to calculate the clinical spinal parameters, vertebral axial rotations, and the 3D centerline of the spinal curvature. A classification based on the centerlines’ axial plane projection was developed. The 3D curvature of the centerline was calculated at each point. A 2D elastic rod finite element model (FEM) of the sagittal spinal curvature for each axial subtype was developed to calculate the 3D deformity of the spine under gravity and axial torsion. Differences in the axial clusters’ clinical parameters, curvature of the spine, and the deformation patterns of the FEM were compared. The characteristics of the sagittal curvature of these axial clusters were determined.

**Results:** Lenke1 was divided into two axial groups (I and II) whereas the Lenke 5 cohort all had the same axial projection pattern (loop shape). T5-T12 kyphosis was significantly different between Lenke1-Group I and the other two groups, p=0.04. The vertebral rotation in both Lenke1-Group I and Lenke 5 had only one maximum value and the FEM deformed in a loop shaped whereas Lenke1-group II showed two maximum values for vertebral rotation and the FEM of the centerline deformed in a lemniscate shape. The ratio of the spinal arc lengths above and below the sagittal inflection points for Lenke1-Groups I and II and Lenke 5 were 0.52, 1.16, and 3.24, respectively.

**Conclusion:** Variations in the axial plane projection of the curve were observed within Lenke types. Lenke 1- Group I and Lenke 5 showed similar 3D curve characteristics suggesting one 3D curve whereas Lenke1-Group II, with two 3D curves, behaved differently. The length of the spinal arcs above and below the sagittal inflection point related to the patterns of axial deformity.

## Introduction

Classification systems of adolescent idiopathic scoliosis (AIS) play an important role in clinical management of the disease [1, 2]. The current widely-used classification system of AIS, Lenke classification, primarily is based on the curve appearance in the frontal plane, with a minor role for the sagittal plane [2]. 3D curve subtypes have been identified within the Lenke subtypes [3, 4]. However, it has not been determined whether the 3D characteristics of the curves follow the current clinical classification of the AIS.

Lenke determined 6 subtypes of AIS with complementary modifiers to provide guidelines for surgical treatment [2]. This system classifies the AIS patients, based on the number of structural curves in the frontal plane to one curve scoliotic types (Lenke1 and Lenke5), two-curve scoliotic types (Lenke 2, 3, and 6), and a three-curve scoliosis (Lenke 4). Three-dimensional classification of the curves resulted in more than 6 curve patterns in AIS, sub classifying the patients within the Lenke types [3–5]. But it was not determined how different aspects of the 3D curves vary within and between the Lenke types. More importantly, it was not determined whether the mechanism of the scoliotic curve development can be explained using the similarity or differences between the 3D curve types. A better description of the fundamental 3D characteristics of the scoliotic curves can have an important role in understanding the pathogenesis of the curve development and subsequent clinical management of the disease [6, 7].

As shown previously, the axial projection of the curve carries important information for surgical and non-surgical management of the right thoracic AIS [6, 8–10]. It was also shown that the sagittal profile strongly relate to the patterns of axial deformation using mechanics of deformation in an elastic rod [3, 11, 12]. Here, it is studied if the axial projection of the curve varies within the Lenke types and whether that variation relates to the sagittal curvature of the spines within these subtypes. As the prevalence of Lenke 1 (thoracic curve) (51%) and Lenke 5 (thoracolumbar/lumbar curve) (12%) is among the highest in AIS patients [13], it was aimed to determine the differences in the 3D curve characteristics of the Lenke 1 and Lenke 5 curves based on an axial classification of the curves. The hypothesis was that variation in the axial plane projections of the curves exists independent of the frontal curve classification but related to the sagittal profile of the spine. The differences between the 3D characteristics of the axial subtypes were used to discuss the mechanics of the various frontal curve pattern developments in these AIS subtypes.

## Methods

### Subjects

A total of 71 Lenke1 and 52 Lenke 5 AIS were selected retrospectively and consecutively. All patients had biplanar standing radiographs and bending films to verify the Lenke types. Male and female patients between 12-17 years old were included. Patients with previous spinal surgery, spondylolisthesis, congenital spinal abnormalities, and neuromuscular conditions were excluded.

### Image processing

3D reconstructions of the spinal radiographs were created in SterEOS 2D/3D (EOS imaging, Paris, France). The 3D reconstruction algorithm allows identifying several anatomical landmarks on the coronal and lateral radiographs to create the 3D morphology of the vertebral bodies based on a statistical model[14]. The 3D models then were used to extract the vertebral centroids and create the 3D centerline of the spine by connecting the 17 vertebrae centroids of the thoracic and lumbar spine [15]. This centerline was isotropically normalized in a way that all the spines have the same heights while modifying the antero-posterior and lateral coordinates by the same factor. The reliability of this method in evaluating the 3D spinal curves was shown previously [3, 15].

### Axial classification of the curves

The 3D centerlines of the Lenke 1 and Lenke 5 patients were clustered based on the variations in the axial projections of the curves. The number of clusters was determined based on the silhouette values [16]. The silhouette value determines the maximum number of the groups with significantly different curve patterns [16]. *K-means* clustering then was performed to divide the cohort into the number of clusters determined by the silhouette value.

### Spinal and pelvic parameters

The spinal and pelvic parameters (Thoracic and lumbar Cobb angles, T5-T12 Kyphosis, L1/S1 lordosis, pelvic incidence, sacral slope, and vertebral axial rotation) were calculated using the 3D models of the spine in each cluster. Vertebral rotation was determined by the angle between the axial projection of the vector connecting the posterior and anterior landmarks on the vertebral endplate and the true horizontal axis (Y-axis) connecting the femoral head centers [17]. The reliability of identifying these landmarks were determined previously [15].

### Geometrical parameter

Using the spinal centerlines, the curvature of the 3D centerline at each vertebral level was calculated. The curvature of the centerline at each point can be considered as the inverse of the radius of a circle that best fits the curvature at that point and mathematically calculates as how rapidly the tangent to the curve changes direction between two consecutive points (Figure 1).

**Figure 1.**
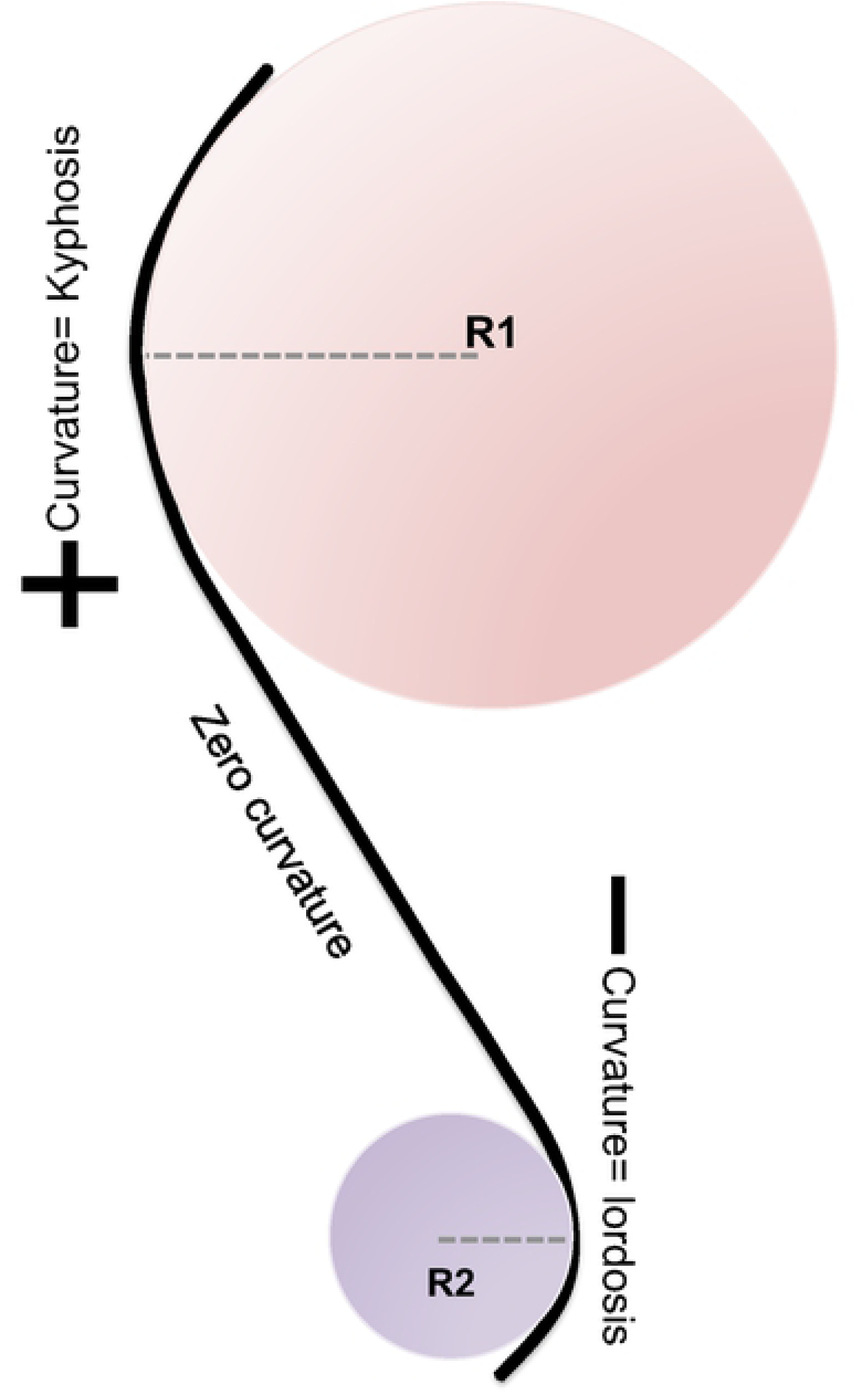
Definition of curvature as the inverse of the radius of a circle (R) that best fits the curve at each point. The positive and negative curvatures and the inflection section (zero curvature) are shown.

### Biomechanical parameters

The sagittal curvature of the spine, without considering the frontal deformity, was used to develop a 2D reduced order finite element model (FEM) of the spine [12]. The S shaped (sagittal profile) 2D model of a homogeneous, isotropic, slender elastic rod (Young modulus=1000Pa and Poisson ratio=0.3) with a circular cross section, was loaded by vertical loading (gravity) at each vertebral level and a torsional moment representing the trunk mass asymmetry as shown in [18]. This model previously generated the 3D deformation patterns of the scoliotic spine [12]. The reduced-order FEM was created for all the axial clusters to calculate the 3D deformation patterns.

Finally, the sagittal curvature of the spine in each axial subtypes were characterized by the length of the positive and negative arcs and the position of the inflection point (Figure 1). The inflection point was determined at the center of the section without any curvature in the sagittal plane (Figure 1).

### Statistical analysis

The clinical, geometrical, and biomechanical parameters were compared between the clusters in Lenke 1 and Lenke 5 cohorts using an analysis of variance followed by a post hoc test (ANOVA and Turkey’s test for normally distributed data and Kruskal-Wallis and Dunn’s test when normality was rejected).

## Results

Silhouette analysis determined two axial clusters in the Lenke 1 cohort: Group I (n=39): loop shaped and Group II (n=32): lemniscate shaped axial projections and one group of axial projection pattern in Lenke 5 (n=52): loop shaped projection. These axial groups and associated frontal and sagittal curves are shown in figure 2. The two most distinguishable axial clusters in Lenke 5 are shown in the supplementary materials (Figure S1). Although both anterior-posterior and lateral deviations of the axial projections were different between the two most distinguished clusters in Lenke 5 patients, the patterns of the axial deformity did not differ between the two clusters (supplementary materials, Figure S1).

**Figure 2.**
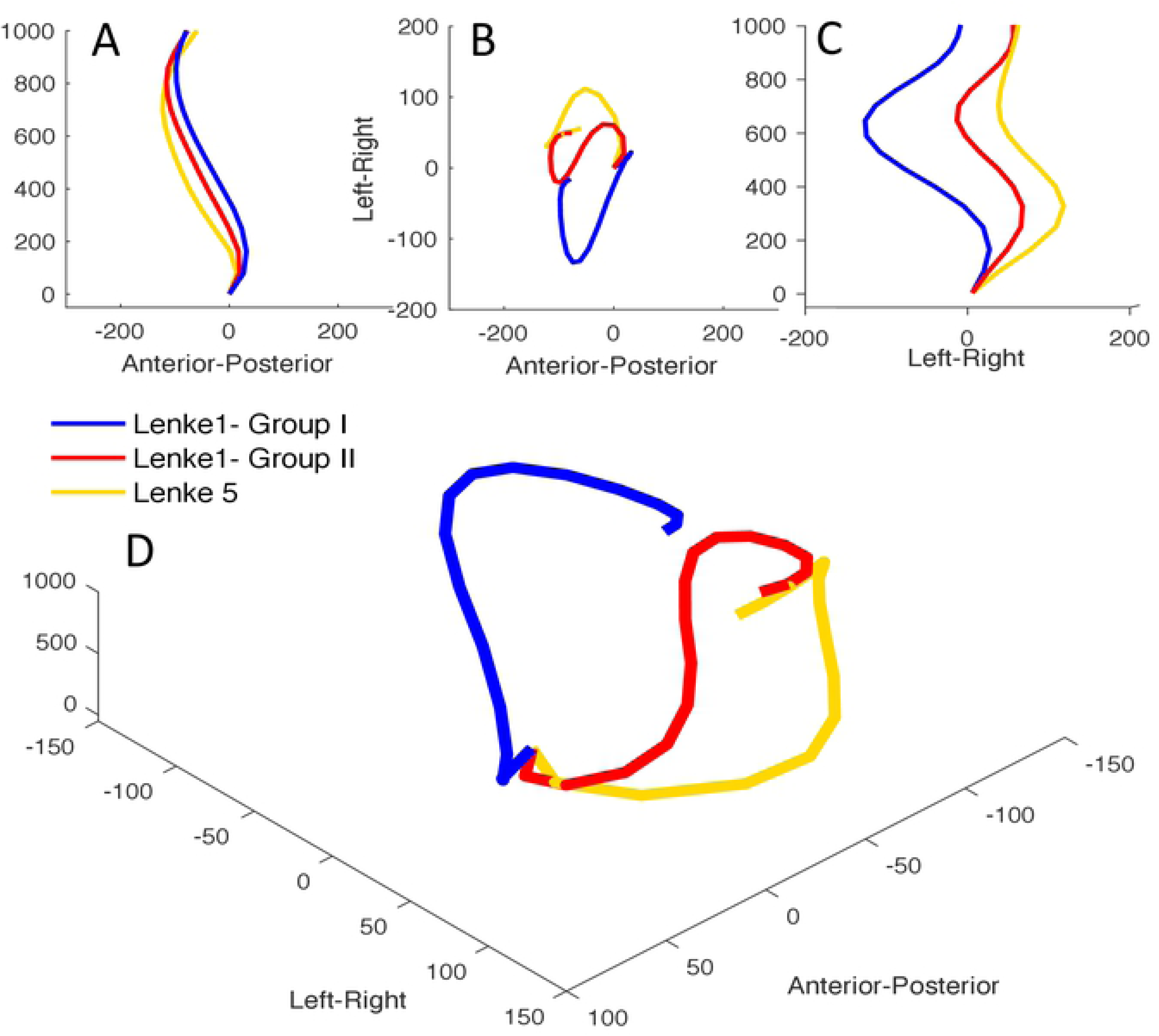
The (A) sagittal, (B) axial, (C) frontal, and (D) 3D spinal centerlines for two subtypes of Lenke 1 (Group I and Group II) and Lenke 5. The X, Y, and Z axes show unite-less normalized distances.

### Spinal and pelvic parameters

Table 1 summarizes the spinal and pelvic measurements in the two clusters of Lenke1 and in Lenke 5. Thoracic Cobb angles were statistically different between the Lenke 1 (Groups I and II) and Lenke 5 cohorts, p=0.032 and p=0.037, respectively. T5-T12 kyphosis was statistically different between Lenke 1-Group I and Lenke 5, p=0.041.

**Table 1.** The spinal and pelvic parameters in the two clusters of the Lenke 1 (Group I and Group II) and Lenke 5.

|  | Thoracic Cobb (°) | Lumbar Cobb (°) | T5-T12 Kyphosis (°) | L1/S1 Lordosis (°) | Pelvic incidence (°) | Sacral slope (°) | Main curve AVR (°) | Main curve AVT (normalized spines) |
| --- | --- | --- | --- | --- | --- | --- | --- | --- |
| Lenke 1- Group I | 60.4±9.3 | 40.2± 5.3 | 14.7±8.7 | 54.5±12.1 | 49.1±10.2 | 43.3± 11.6 | 13.8±6.4 | 0.2±0.04 |
| Lenke 1- Group II | 56.6±10.1 | 42.9±7.5 | 20.6±9.5 | 55.3±14.2 | 52.6±9.8 | 44.5±10.0 | 16.2±8.8 | 0.05±0.00 |
| Lenke 5 | 25.7 ±4.2 | 43.5±12.5 | 22.6±9.7 | 51.8±10.2 | 55.5±9.3 | 40.8±12.6 | 27.2±8.5 | 0.15±0.01 |

Figure 3 shows the axial rotation of the vertebral bodies in the three clusters. Similar patterns were observed for Lenke 1- Group I and Lenke 5 patients with one peak (maximal) for vertebral rotation whereas the vertebral rotation plot for Lenke 1- Group II showed two peaks related to the axial rotation of the apical vertebrae in the thoracic and lumbar curves (Figure 3). This suggests Lenke 1-Group I and Lenke 5 are the true “single curves” whereas Lenke1-Group II has two 3D curves, without the lumbar being structural yet.

**Figure 3.**
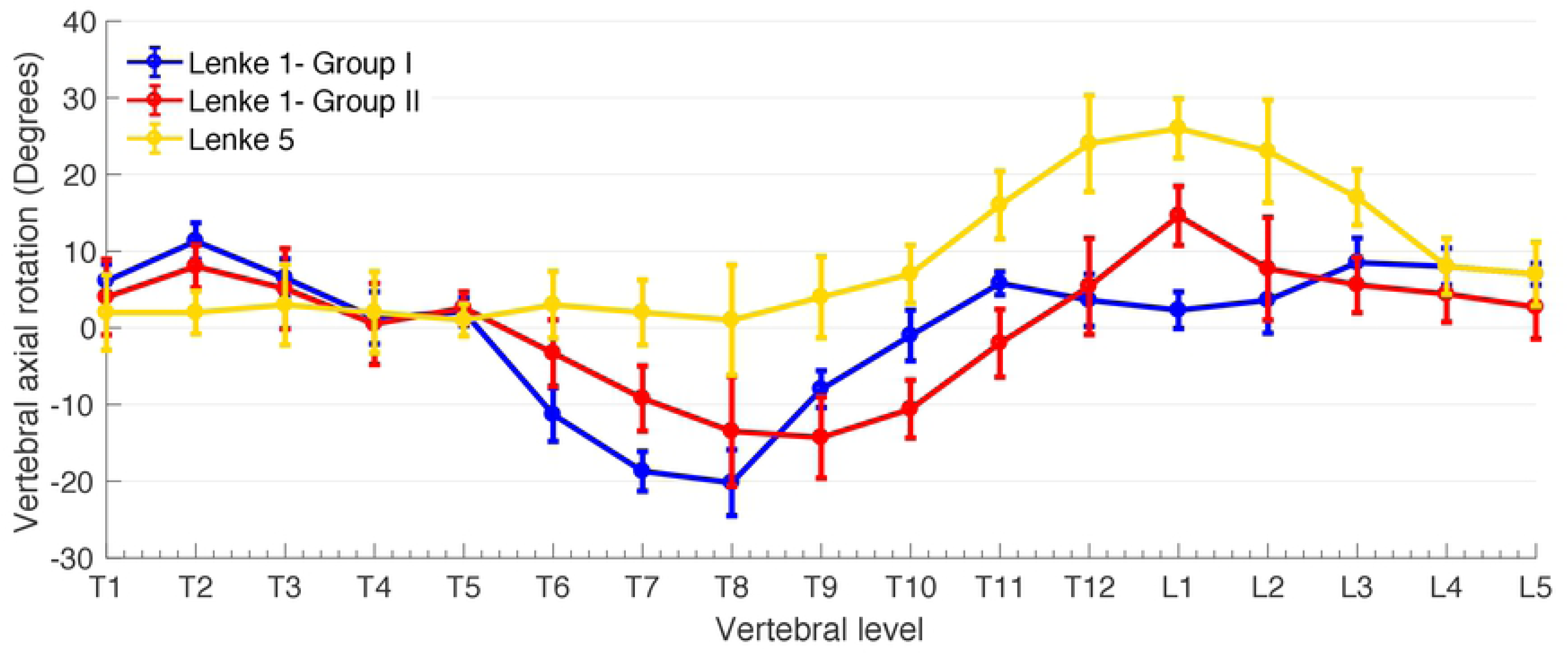
The vertebral axial rotation in Lenke 1-Group I, Lenke 1-Group II, and Lenke 5. The vertebral rotations present with two peaks for Lenke1-Group II patients while the other two groups show only one maximum vertebral rotation.

### Geometrical parameters

Figure 4 shows the curvature vector, representing its magnitude, of the three curves at each vertebral level along the spine in three views (Figure 4A) and in 3D (Figure 4B). The curvature of the spine changes rapidly in Lenke 1- Group II whereas the curvature in Lenke1- Group I and Lenke 5 remained relatively constant along the spine (Figure 4C). The 3D curvatures of the centerline in Lenke1- Group I and Lenke 5 were close to zero along the spine ranging between [-0.001, 0.001] (Figure 4C). On the other hand, in Lenke 1-Group II, the curvature along the spine changed direction with values exceeding the curvature in the other two subtypes (Figure 4C.)

**Figure 4.**
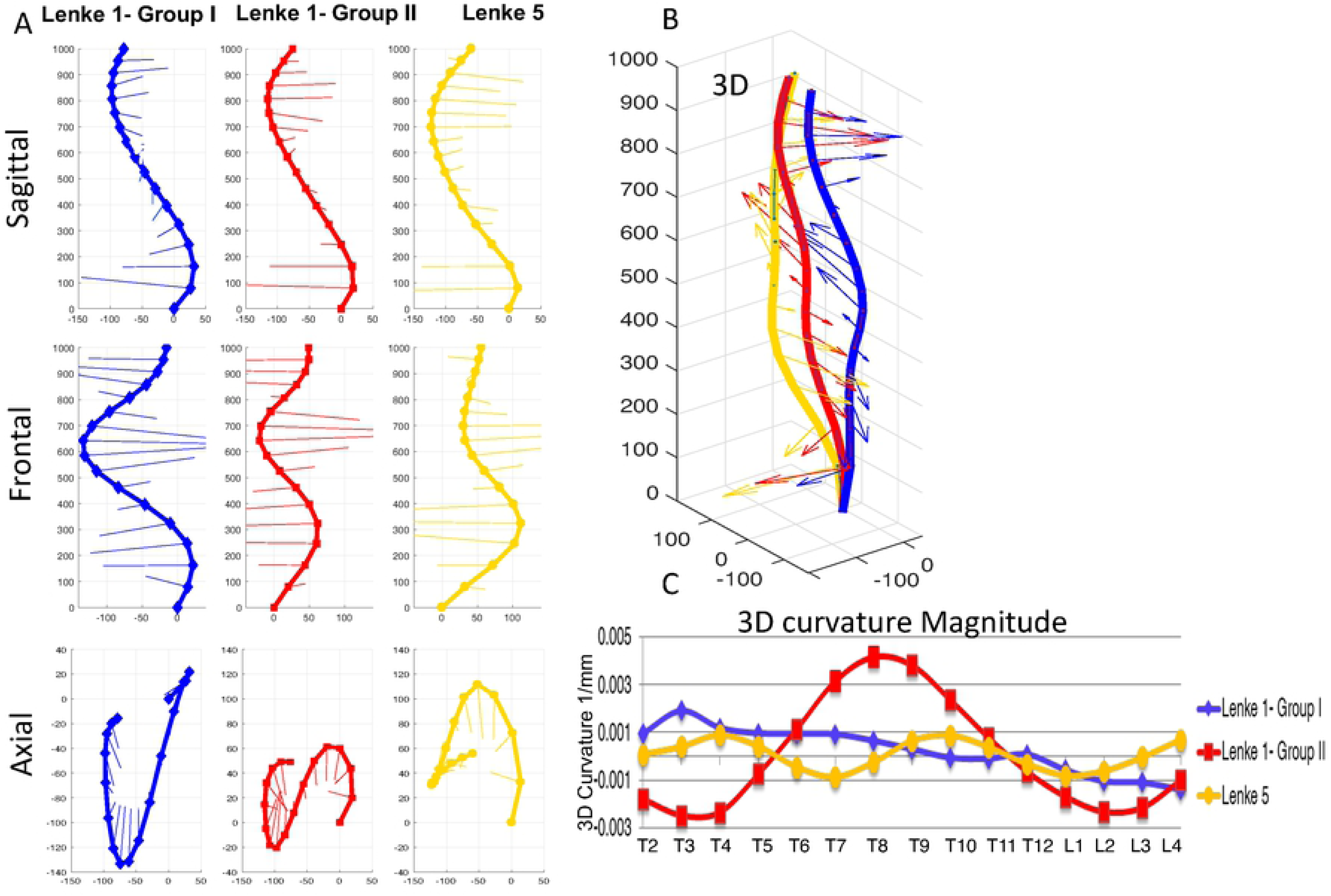
A) The curvatures of the spine in the sagittal, frontal and axial plane are presented in the three groups. The positive curvature in the sagittal view determines the kyphotic section of the spine (positive curvature) and the negative curvature determines the lordotic section of the spine (negative curvature). The point or section of the spine without any curvature is the inflection point/section B) The 3D curvature of the three groups. C) The magnitudes of the 3D curvature for the three subtypes are shown. Lenke1-Group II shows three peaks for the 3D curvature while the curvature in Lenke 1-Group I and Lenke 5 remains close to zero.

### Sagittal variation in the curves

The inflection points of the sagittal curves were at T7 for Lenke1- Group I, T11 Lenke1- Group II, and L2 in Lenke 5 patients (Figure 4A). The ratio of the arc length above and below the inflection point was 0.52, 1.16, and 3.24, for Lenke1- Groups I and II and Lenke 5 respectively.

### Biomechanical parameters

Figure 5 shows the deformation patterns of the 2D finite element models, of the S shaped sagittal geometry as shown in figure 2A. The deformation patterns of the Lenke 1- Group I and Lenke 5 were looped shaped whereas the Lenke 1-Group II showed a lemniscate axial projection, as also was seen in the clinical data (Figure2 B).

**Figure 5.**
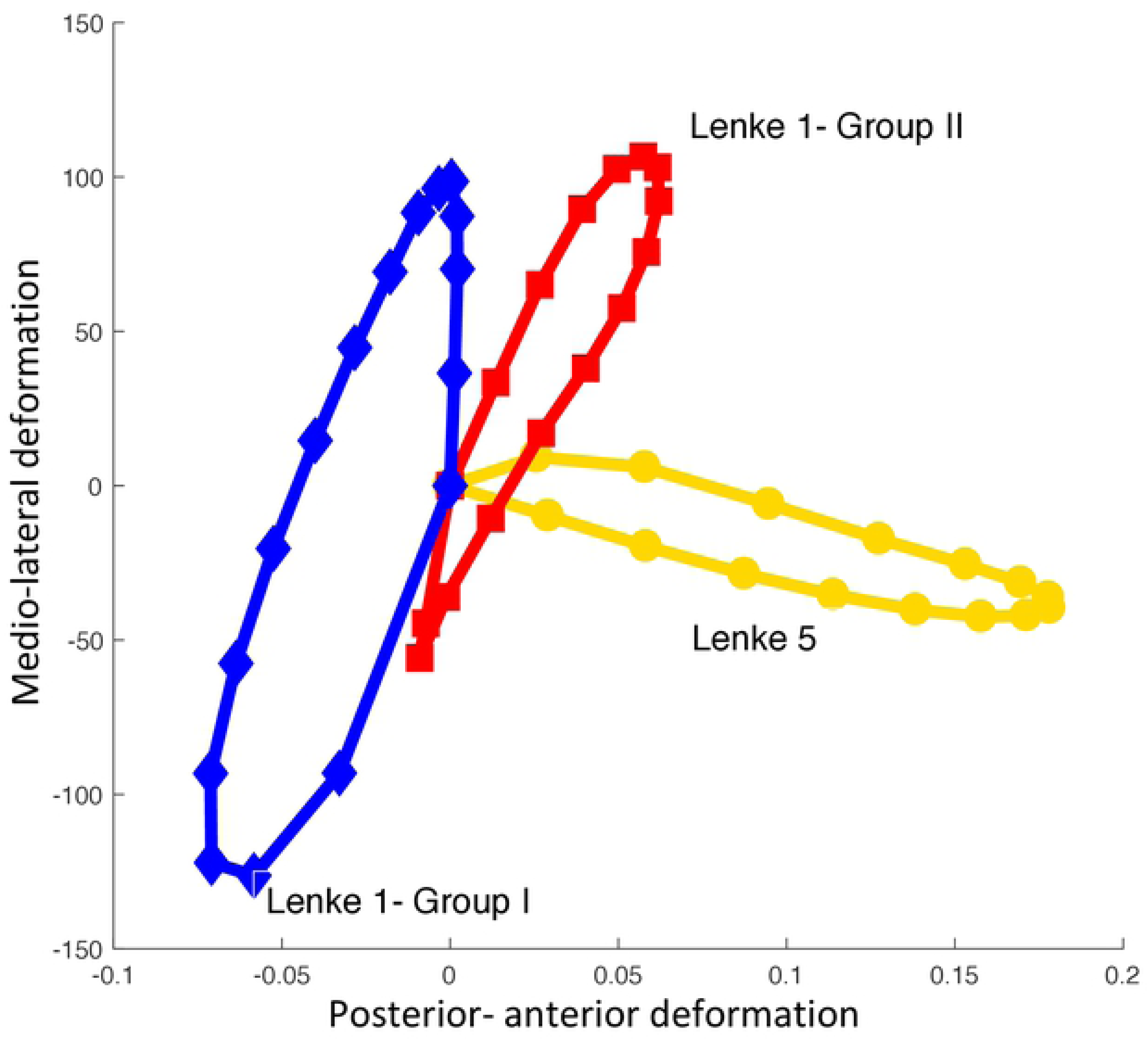
The finite element deformations of the three subtypes. Lenke1-Group I and Lenke 5 projection have a loop shaped axial projection, although in different directions. Lenke1-Group II formed a lemniscate shaped axial projection. The X and Y-axes show unite-less normalized distances.

## Discussion

This study aimed to show how the intrinsic characteristics of the 3D scoliotic curves vary within and between the two most prevalent AIS curve types *i.e.,* Lenke1 and Lenke 5. Our results determined that a subgroup of Lenke 1 (Group I) and Lenke 5 patients present with similar shape of axial projection (loop shape) showing one true 3D curve whereas the Lenke 1-Group II showed two 3D curves and a lemniscate axial projection (Figure 6). Comparing the sagittal curvature of these subtypes revealed that the length of the positive and negative curvatures of the sagittal profile was significantly different in Lenke1- Group I and Lenke 5 whereas the lengths of these two arcs were more similar, compared to the other two groups, in Lenke1-Group II (Figure 6). This result showed the link between the axial projection, sagittal profile, and the coronal deformity of the spine in scoliotic subtypes.

**Figure 6.**
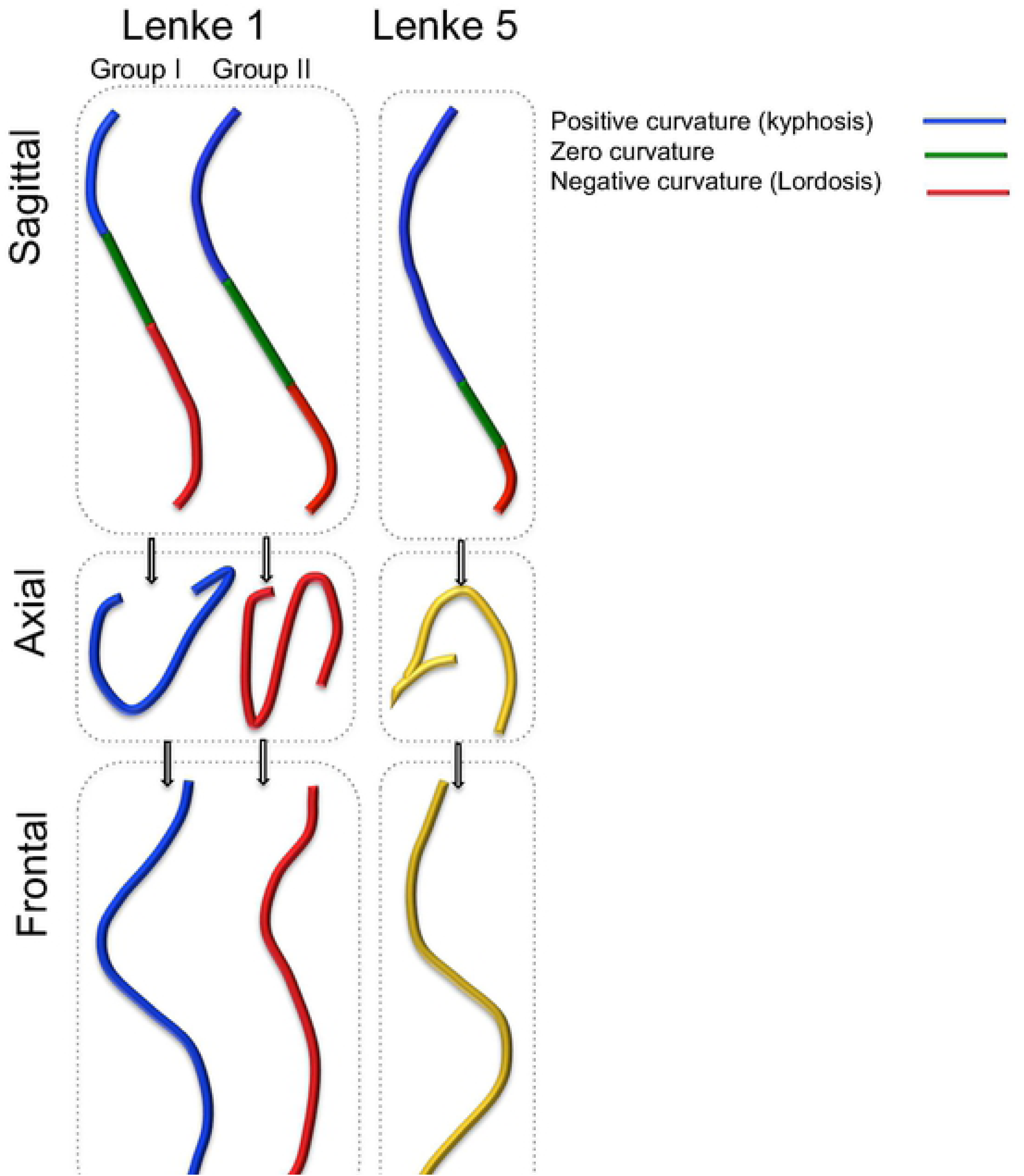
Schematics of the sagittal curvature and the corresponding axial and frontal curve deformation in Lenke 1 and Lenke 5 AIS subtypes. The positive curvature (kyphotic), negative curvature (lordotic), and the section without curvature are shown in the sagittal view.

It was previously showed that two axial subgroups exist in right thoracic AIS: loops shaped and lemniscate shaped [3, 19]. The sagittal curvature of the spine relates to the pattern of the 3D deformity development, particularly the aforementioned axial subgroups of right thoracic AIS [19]. When the spine was considered as a curved elastic rod, it was shown quantitatively that the distribution of the moments along the spine are altered by the initial geometry of the sagittal spinal curvature which in turn results in different 3D deformation patterns among scoliotic patients [11, 19]. Considering the sagittal profiles in the subgroups of Lenke1 (Figure 6), it is expected that the two sagittal subtypes deform differently. The sagittal profile of the spine is characterized in Lenke 1-Group I with one large lordotic segment (negative curvature) with few vertebrae in the kyphotic section of the spine (Figure 6). The sagittal profile of the spine in Lenke1- Group II, on the other hand, has two curves where the lengths of the positive and negative curvature arcs are closer in magnitude (Figure 6). Lenke 5 presents with one large kyphotic section (positive curvature) with few vertebrae in the lordotic section of the spine (Figure 6). As the moments that rotate the spine off the sagittal plane are a function of the sagittal curvature, particularly the angle between the tangent to the spine and horizontal line [11], it is expected that the maximum off-plane deformation occurs at different locations of the spine in these subtypes (Figure 7A). The area above the lower apex in the sagittal plane in Lenke 5 and the area below the upper apex in Lenke 1- Group I are subject to maximum moment due to the inclination of the curves in those regions. These moments that bend the curve off the sagittal plane are responsible for frontal deformity in the thoracolumbar/ lumbar section in Lenke 5 and in the mid thoracic section in Lenke 1-Group 1 (figure 7B). As a result, the deformed spines, in both Lenke1- Group I and Lenke 5 when projected on to the axial plane form a loop shape, *i.e.,* only one 3D curve. In Lenke 1- Group II the sections with minimal tangent angle (maximum moment) fall between the two apices of the sagittal curve resulting in deformation of the kyphotic curve above the inflection point and deformation of the lordotic curve below the inflections point, creating two 3D curves in this subtypes. These two curves when projected on the axial plane form a lemniscate shape (Figure 7). This explains why the axial plane curve characteristics are similar in Lenke1-Group I and Lenke 5 but differ from Lenke1-Group II.

**Figure 7.**
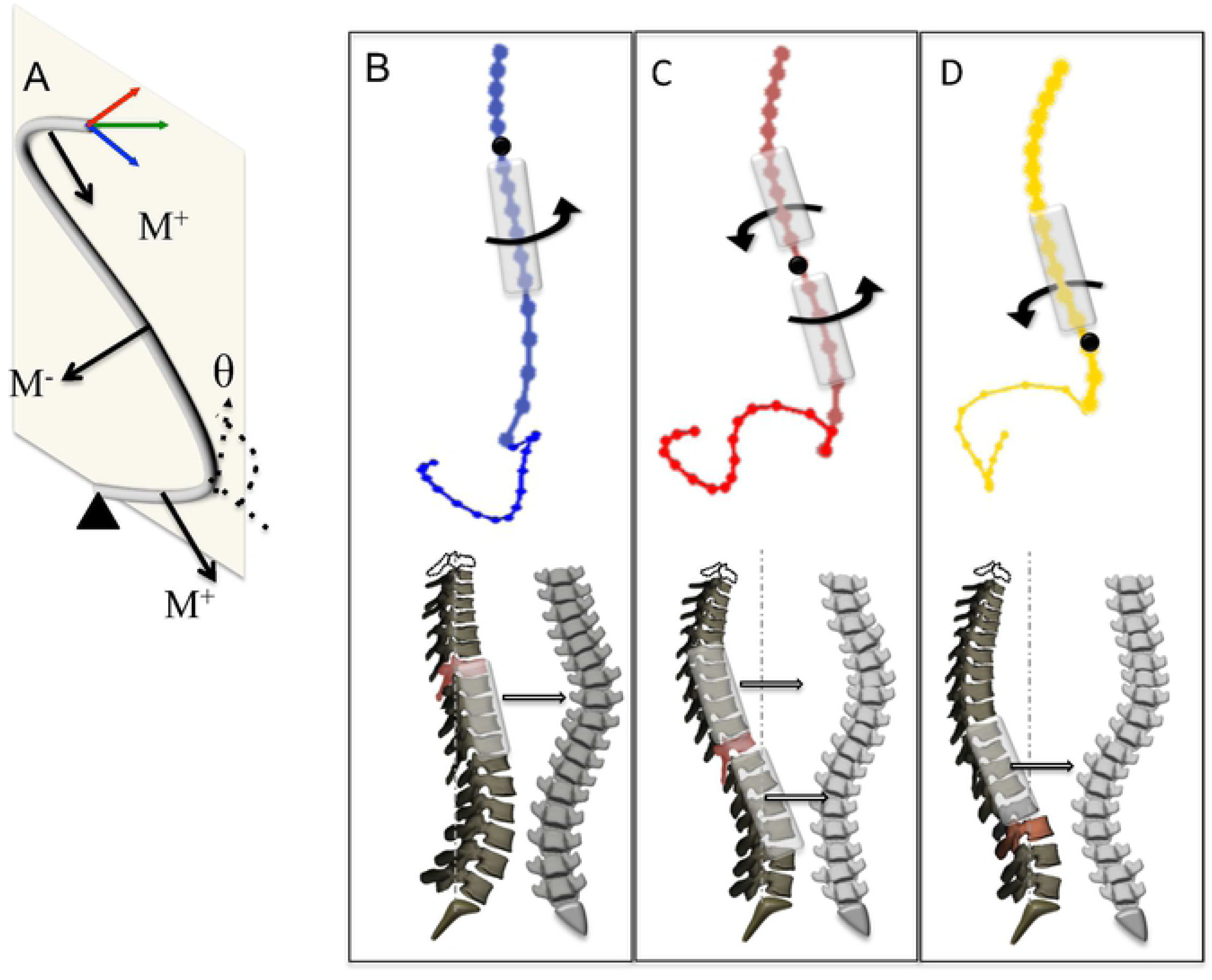
Mechanism of the curve development in the three subtypes. A) The sagittal curvature of the spine (tangent angle to the curve, θ) determines the direction of the moments that deflects the curve off the sagittal plane. B) In Lenke 1-Group I, with a long lordotic curve, the maximum moment is caudal to the inflection point and above the lordosis apex in the sagittal plane thus the maximum deformity occurs in main thoracic (Shaded section). C) In Lenke 1-Group II, with kyphotic and lordotic curve close in length, the areas between the inflection points and the two apices in the sagittal plane is prone to off-plane deformity (shaded sections) D) In Lenke 5, the kyphotic area above the inflection point and below the kyphotic apex in the sagittal plane deforms off plane (shaded section) resulting in a thoracolumbar/lumbar frontal deformity. It should be noted that the location of frontal curve is not at the same level as the 3D curve as shown in figure 1.

The mechanics of elastic rods were used to explain scoliotic curve development in AIS from a mechanical standpoint [11, 12]. It was shown that higher rod inclination results in an increased moment on the rod [11]. This theory is similar in concept to the posterior shear theory that believes an increased posterior shear in posteriorly inclined vertebrae results in rotational instability of the spine[20]. However, different from the shear theory, that was based on the preexisting rotation of the vertebral bodies [21, 22], the rod theory showed that physiological loading of the spine can result in rotation of the kyphotic and lordotic sections of a 2D S shaped curve while exposing the most inclined vertebrae (posteriorly or anteriorly) to higher 3D deformations (Figure 7). The rod theory also differs from the shear theory, as in the latter the shear force is responsible for creating a moment that rotates the vertebrae whereas the rod theory explain that the preexisting rotation in the spine is a results of the rotational moments on the spine which if cannot be tolerated by the stiffness of the spine result in excessive 3D deformation. In other words, the rod theory explain the rotational instability of the spine as a result of the physiological loading while the shear theory use an already rotated spine to underline the role of the posterior shear on rotational instability. Other way said, the shear theory does not explain the preexisting rotation of the spine while the rod theory does. The differentiation between the mechanism of the curve development is important in prevention of the curve development because is shows that scoliosis initiates with an off-plane deflection of the 2D sagittal curves at the sections with maximum inclination (posteriorly or anteriorly) as opposed to rotational instability as a result of shear load or retrolisthesis in those sections.

The role of the spinal slenderness [23] and flexibility of the spine cannot be ignored in explaining this deformation phenomenon and induction of scoliosis. Based on the rod theory of scoliotic development, both anteriorly and posteriorly tilted vertebrae may result in increased moments and if the section with such alignment is long or alternatively flexible enough, the acting moment overcome the stiffness of the curve and will deflect the spine off the sagittal plane and result in a 3D deformity. The forward inclined vertebrae are close to the caudal and cranial ends of the spine, with a shorter length compared to the section of the spine between the two sagittal apices (Figure 6A and Figure 7). It is expected that excessive forward tilt of the vertebrae above and below the cranial and caudal apices also result in the curve deflection, given that the moment is large enough to exhaust the bending modulus of the spine. If the two apices of the sagittal curve are close to each other, the anteriorly inclined vertebrae above or below the cranial and caudal curves can also rotate and create a third 3D curve as seen in Lenke 4 or compensatory curve in other curve types. Finally, while studies have alluded to the genetic factors affecting the morphology of the hip and knee and the pathway of developmental musculoskeletal diseases[24, 25], the role of genetics in variation is sagittal alignment of the spine remains to be further explored.

The clinical classification of the spine should aim to guide the surgical decision-making. In the context of selective fusion of the spine in AIS, considering the described 3D characteristics of the curve in this study can guide the fusion level selection. A short fusion that does not include the entire 3D curve in subjects with a loop shaped axial projection (Lenke 1-Group I and Lenke 5) can lead to postural compensation and need for revision surgery[26]. In Lenke 1- Group II with two 3D curves, and the secondary curve not being structural yet, it is expected that fusion of the main thoracic curve, as an adequate kyphosis is being imparted, can change the force distribution along the spine and de-twist the secondary curvature (lumbar) spontaneously. Finally, it is expected that the Lumbar modifier As and hypokyphosis Lenek1s are mostly in Lenke1- Group I and lumbar modifier Cs and normal/hyperkyphosis in Lenke1-Group II[3], however considering the true number of the 3D curves can eliminate the error associated with the classification of the borderline cases and improve the surgical planning.

In summary, as the scoliosis impacts the spinal alignment in the three anatomical planes, understanding the 3D characteristic of the scoliotic curves is key for systematic classification of the patients both for etiological studies and treatment of the patients. The mechanism of the curve development in AIS, for different frontal curve pattern, was explained. The 3D curve patterns are dictated by the length of the spine with positive and negative curvatures (figure 1) and the position of the frontal curve is dictated by the position of the most tilted vertebrae with respect to the sagittal apices of the curve (Figure 6 and 7). Consequently, this explanation of AIS curve development leads us to determining the mechanical risk factors of the scoliotic curve development in pediatric population: 1- Flexible spine. 2- Slender spine-narrow cross section with respect to the length- (this condition can be satisfied as a result of fast spinal growth at puberty). 3- the relative length of the sagittal spinal arcs with positive and negative curvature (as shown to determine the number of 3D curves). 4- long section with tilted vertebrae (forward or backward) in the sagittal plane (as shown to determine the location of the frontal curve).

## Acknowledgment

I acknowledge professors Rene Castelein and Prashant Purohit for insightful discussions.

## References

1. Lenke LG, Betz RR, Haher TR, Lapp MA, Merola AA, Harms J, Shufflebarger HL (2001) Multisurgeon assessment of surgical decision-making in adolescent idiopathic scoliosis: curve classification, operative approach, and fusion levels. Spine (Phila Pa 1976) 26:2347–2353

2. Lenke LG, Betz RR, Harms J, Bridwell KH, Clements DH, Lowe TG, Blanke K (2001) Adolescent idiopathic scoliosis: a new classification to determine extent of spinal arthrodesis. J Bone Joint Surg Am 83-A:1169–1181

3. Pasha S, Hassanzadeh P, Ecker M, Ho V (2019) A hierarchical classification of adolescent idiopathic scoliosis: Identifying the distinguishing features in 3D spinal deformities. PLoS One 14:e0213406. doi: 10.1371/journal.pone.0213406

4. Duong L, Mac-Thiong JM, Cheriet F, Labelle H (2009) Three-dimensional subclassification of Lenke type 1 scoliotic curves. J Spinal Disord Tech 22:135–143. doi: 10.1097/BSD.0b013e31816845bc

5. Kadoury S, Labelle H (2012) Classification of three-dimensional thoracic deformities in adolescent idiopathic scoliosis from a multivariate analysis. Eur Spine J 21:40–49. doi: 10.1007/s00586-011-2004-2

6. Pasha S, Baldwin K (2019) Surgical outcome differences between the 3D subtypes of right thoracic adolescent idiopathic scoliosis. Eur Spine J 28:3076–3084. doi: 10.1007/s00586-019-06145-4

7. Shen J, Kadoury S, Labelle H, Parent S (2016) Geometric Torsion in Adolescent Idiopathic Scoliosis: A Surgical Outcomes Study of Lenke Type 1 Patients. Spine (Phila Pa 1976) 41:1903–1907. doi: 10.1097/BRS.0000000000001651

8. Pasha S (2019) 3D spinal and rib cage predictors of brace effectiveness in adolescent idiopathic scoliosis. BMC Musculoskelet Disord 20:384. doi: 10.1186/s12891-019-2754-2

9. Pasha S, Cahill PJ, Flynn JM, Sponseller P, Newton PO, Group aHS (2018) Relationships Between the Axial Derotation of the Lower Instrumented Vertebra and Uninstrumented Lumbar Curve Correction: Radiographic Outcome in Lenke 1 Adolescent Idiopathic Scoliosis With a Minimum 2-Year Follow-up. J Pediatr Orthop 38:e194–e201. doi: 10.1097/BPO.0000000000001136

10. Almansour H, Pepke W, Bruckner T, Diebo BG, Akbar M (2019) Three-Dimensional Analysis of Initial Brace Correction in the Setting of Adolescent Idiopathic Scoliosis. J Clin Med 8. doi: 10.3390/jcm8111804

11. Purohit P, Pasha S (2020) A semi-analytical model of pediatric spine. In. J. Biomech.

12. . Pasha S (2019) 3D Deformation Patterns of S Shaped Elastic Rods as a Pathogenesis Model for Spinal Deformity in Adolescent Idiopathic Scoliosis In. Scientific Reports.

13. Lenke LG, Betz RR, Clements D, Merola A, Haher T, Lowe T, Newton P, Bridwell KH, Blanke K (2002) Curve prevalence of a new classification of operative adolescent idiopathic scoliosis: does classification correlate with treatment? Spine (Phila Pa 1976) 27:604–611. doi: 10.1097/00007632-200203150-00008

14. Humbert L, De Guise JA, Aubert B, Godbout B, Skalli W (2009) 3D reconstruction of the spine from biplanar X-rays using parametric models based on transversal and longitudinal inferences. Med Eng Phys 31:681–687. doi: 10.1016/j.medengphy.2009.01.003

15. Pasha S, Schlosser T, Zhu X, Mellor X, Castelein R, Flynn J (2017) Application of Low-dose Stereoradiography in In Vivo Vertebral Morphologic Measurements: Comparison With Computed Tomography. J Pediatr Orthop. doi: 10.1097/BPO.0000000000001043

16. Kaufman L, P.J. R (1990) Finding Groups in Data: An Introduction to Cluster Analysis

17. Illes T, Somoskeoy S (2013) Comparison of scoliosis measurements based on three-dimensional vertebra vectors and conventional two-dimensional measurements: advantages in evaluation of prognosis and surgical results. Eur Spine J 22:1255–1263. doi: 10.1007/s00586-012-2651-y

18. de Reuver S, Brink RC, Homans JF, Kruyt MC, van Stralen M, Schlosser TPC, Castelein RM (2019) The Changing Position of the Center of Mass of the Thorax During Growth in Relation to Pre-existent Vertebral Rotation. Spine (Phila Pa 1976) 44:679–684. doi: 10.1097/BRS.0000000000002927

19. Pasha S (2019) 3D Deformation Patterns of S Shaped Elastic Rods as a Pathogenesis Model for Spinal Deformity in Adolescent Idiopathic Scoliosis. Sci Rep 9:16485. doi: 10.1038/s41598-019-53068-7

20. Schlösser TP, Shah SA, Reichard SJ, Rogers K, Vincken KL, Castelein RM (2014) Differences in early sagittal plane alignment between thoracic and lumbar adolescent idiopathic scoliosis. Spine J 14:282–290. doi: 10.1016/j.spinee.2013.08.059

21. Janssen MM, Kouwenhoven JW, Schlösser TP, Viergever MA, Bartels LW, Castelein RM, Vincken KL (2011) Analysis of preexistent vertebral rotation in the normal infantile, juvenile, and adolescent spine. Spine (Phila Pa 1976) 36:E486–491. doi: 10.1097/BRS.0b013e3181f468cc

22. Kouwenhoven JW, Smit TH, van der Veen AJ, Kingma I, van Dieën JH, Castelein RM (2007) Effects of dorsal versus ventral shear loads on the rotational stability of the thoracic spine: a biomechanical porcine and human cadaveric study. Spine (Phila Pa 1976) 32:2545–2550. doi: 10.1097/BRS.0b013e318158cd86

23. Chen H, Schlösser TPC, Brink RC, Colo D, van Stralen M, Shi L, Chu WCW, Heng PA, Castelein RM, Cheng JCY (2017) The Height-Width-Depth Ratios of the Intervertebral Discs and Vertebral Bodies in Adolescent Idiopathic Scoliosis vs Controls in a Chinese Population. Sci Rep 7:46448. doi: 10.1038/srep46448

24. Baird DA, Evans DS, Kamanu FK, Gregory JS, Saunders FR, Giuraniuc CV, Barr RJ, Aspden RM, Jenkins D, Kiel DP, Orwoll ES, Cummings SR, Lane NE, Mullin BH, Williams FM, Richards JB, Wilson SG, Spector TD, Faber BG, Lawlor DA, Grundberg E, Ohlsson C, Pettersson-Kymmer U, Capellini TD, Richard D, Beck TJ, Evans DM, Paternoster L, Karasik D, Tobias JH (2019) Identification of Novel Loci Associated With Hip Shape: A Meta-Analysis of Genomewide Association Studies. J Bone Miner Res 34:241–251. doi: 10.1002/jbmr.3605

25. Pregizer SK, Kiapour AM, Young M, Chen H, Schoor M, Liu Z, Cao J, Rosen V, Capellini TD (2018) Impact of broad regulatory regions on Gdf5 expression and function in knee development and susceptibility to osteoarthritis. Ann Rheum Dis 77:450. doi: 10.1136/annrheumdis-2017-212475

26. Pasha S, Baldwin K (2019) Surgical outcome differences between the 3D subtypes of right thoracic adolescent idiopathic scoliosis. Eur Spine J. doi: 10.1007/s00586-019-06145-4

27. de Reuver S, Brink RC, Homans JF, Kruyt MC, van Stralen M, Schlösser TPC, Castelein RM (2019) The Changing Position of the Center of Mass of the Thorax During Growth in Relation to Pre-existent Vertebral Rotation. Spine (Phila Pa 1976) 44:679–684. doi: 10.1097/BRS.0000000000002927

